# Comparing Electrical and Ultrasound Transcutaneous Vagus Nerve Stimulation (taVNS) on Associative Memory

**DOI:** 10.64898/2026.02.23.707405

**Authors:** Benjamin J. Griffiths, Ziqi He, Irem Ciftepinar, Hyuk Choi, Jae-Jun Song, Marcus Kaiser, JeYoung Jung

## Abstract

Associative memory, the ability to bind and retrieve relationships between unrelated elements, is a cornerstone of human cognition and a primary target for neurorehabilitation. Vagus nerve stimulation (VNS) has emerged as a promising method to modulate the locus coeruleus-norepinephrine (LC-NE) system and hippocampal-prefrontal circuits essential for memory. However, the comparative efficacy of non-invasive modalities such as electrical (E-taVNS) and the emerging field of ultrasound (U-taVNS) remains poorly understood in the context of active recall. In this study, participants performed a crossmodal video–word associative memory task before and after receiving either E-taVNS or U-taVNS in active and sham conditions. We investigated whether these modalities enhance cued recall accuracy and retrieval reaction time. Our results revealed that neither E-taVNS nor U-taVNS significantly improved recall accuracy. However, E-taVNS significantly accelerated response times specifically for correctly recalled items. These findings suggest that while taVNS may not increase the likelihood of recalling associative memories, electrical stimulation may enhances the efficiency in which we do so. These findings suggest that electrical taVNS is a viable tool for facilitating memory search processes, though further research is required to optimize ultrasound parameters and validate mechanistic pathways through physiological monitoring.

## Introduction

Memory enables us to encode, store, and retrieve past experiences and forms a core foundation of human cognition (Squire 2009). Among different memory systems, associative memory is the ability to bind and retrieve relationships between otherwise unrelated elements and is critical for daily learning, decision-making, and constructing coherent situational representations (Tulving and Thomson 1973; Davachi 2006). Extensive evidence suggests the hippocampus and broader medial temporal lobe (MTL) structures as central to associative binding, integrating multimodal information into unified memory traces (Konkel and Cohen 2009).

Associative memory also depends strongly on prefrontal executive control, particularly during strategic encoding or selective retrieval (Eichenbaum 2017). Lifespan neuroimaging studies show that frontal grey matter volume predicts associative performance even more robustly than MTL structures, highlighting the role of prefrontal–hippocampal interactions (Guardia et al. 2023). As associative memory relies heavily on hippocampal integrity, it is particularly vulnerable to age-related decline and is recognised as an early cognitive marker of Alzheimer’s disease (Algarabel et al. 2012; Bastin et al. 2014). Its dependence on neuromodulatory systems, especially norepinephrine (NE), further positions associative memory as a compelling target for interventions that modulate hippocampal–prefrontal networks (Mayes et al. 2007; Mather et al. 2016).

One promising intervention is vagus nerve stimulation (VNS). The vagus nerve (VN) is a key component of the parasympathetic nervous system and consists primarily of afferent fibres (∼80%) projecting from the periphery to central neuromodulatory hubs, including the nucleus tractus solitarius (NTS), locus coeruleus (LC), and basal forebrain (Berthoud and Neuhuber 2000; Bonaz et al. 2017). Through these pathways, vagal activation modulates neurotransmitter systems, most prominently NE and acetylcholine, linked to attention, emotional processing, and memory (Vonck et al. 2014). Although direct VNS (dVNS) has demonstrated therapeutic efficacy for epilepsy and depression (Ben-Menachem 2002; George et al. 2003; Carreno and Frazer 2017), its invasive nature limits feasibility for cognitive research in healthy populations.

Growing recognition of the therapeutic potential of vagal modulation has stimulated increasing interest in non-invasive alternatives, most notably transcutaneous vagus nerve stimulation (taVNS). Two principal modalities have emerged: electrical VNS (eVNS) and, more recently, ultrasound-based VNS (uVNS). eVNS is typically delivered auricularly at the cymba conchae of both ears using surface electrodes to stimulate afferent vagal fibres, thereby modulating LC–hippocampal and prefrontal circuits and providing a non-invasive route to influencing cognitive processes (Frangos et al. 2015; Yakunina et al. 2017). uVNS involves the external application of focused ultrasound pulses to the auricular branch of the vagus nerve, allowing deeper energy penetration and potentially more selective engagement of vagal fibres (Zhang et al. 2025).

A few behavioural studies have explored whether taVNS can enhance memory performance (Naparstek et al. 2023). Early findings showed that a single session of taVNS improved face–name associative recall in older adults (Jacobs et al. 2015) and enhanced verbal order memory, but only for phonologically similar items (Kaan et al. 2021). These results align with broader evidence suggesting that eVNS can enhance memory (Colzato and Beste 2020; Wang et al. 2024). Neuroimaging and neurophysiological studies provide a mechanistic basis for these effects: : eVNS modulates brainstem and limbic circuits, decreasing BOLD signals in the LC, hippocampus, and parahippocampal gyrus, while increasing activation in the insula, anterior cingulate, inferior and superior frontal gyri, caudate, and putamen (Kraus et al. 2007; Rajiah et al. 2024). Recent work also shows that taVNS increases hippocampal connectivity with the prefrontal cortex and cingulate cortex in individuals with MCI and AD (Murphy et al. 2023). Combined with EEG, a study reported that taVNS enhanced P3b amplitudes (Ventura-Bort et al. 2018) and another study showed that taVNS attenuated alpha oscillations power related to the LC-NE system (Sharon et al. 2021). However, behavioural findings remain inconsistent: some studies report no memory improvement in young adults (Mertens et al. 2020), and evidence for uVNS is largely preclinical (Imamura et al. 2023), with no human memory studies to date. Thus, despite strong theoretical and neurobiological rationale for LC–NE-driven modulation of hippocampal–prefrontal circuits, the reliability of tVNS-related associative memory enhancement remains unclear.

Here, we investigated the effects of both eVNS and uVNS on associative memory. Participants completed a video–word associative memory task before and after taVNS stimulation and, prior to the experiment, filled out questionnaires assessing anxiety, depression, and interoceptive perception. Each participant received either E-taVNS or U-taVNS in both active and sham conditions. We hypothesised that active E-taVNS and U-taVNS would enhance associative memory performance relative to both pre-stimulation and sham. Additionally, we directly compared the efficacy of E-taVNS and U-taVNS and examined whether baseline traits predicted individual variability in taVNS-induced memory changes.

## Methods and Materials

### Participants

An a priori power analysis was conducted using G*Power 3.1.9.7 (Faul et al. 2009) to estimate the required sample size for a within-subjects design (power = .80, α = .05). Assuming a moderate effect size based on previous findings (Johnson & Steenbergen, 2022), the minimum required sample was 19 participants. To ensure adequate statistical power and allow for possible attrition, we aimed to recruit 30 participants per stimulation modality.

A total of 59 healthy adults (46 females, 13 males; M = 23.6 years, SD = 2.88) took part in the study. Participants were randomly allocated to either the electrical stimulation group (eVNS; n = 30) or the ultrasound stimulation group (uVNS; n = 29).

All volunteers completed a VNS Safety Questionnaire to confirm eligibility. Exclusion criteria included: age below 18; presence of any active implanted medical device (e.g., pacemaker, cochlear implant); history of carotid atherosclerosis or cervical vagotomy; cardiovascular abnormalities (hypertension, hypotension, bradycardia, tachycardia); metallic implants in the head or neck; current or past neurological or psychiatric disorders; fainting tendency; family history of epilepsy; current use of psychoactive medication (except hormonal contraception); and pregnancy.

All participants provided written informed consent for participation and publication of anonymised results. The study received ethical approval from the University of Nottingham Ethics Committee (F1619R) and adhered to the Declaration of Helsinki.

### Transcutaneous auricular vagus nerve stimulation (taVNS)

taVNS was delivered using two modalities: electrical stimulation (Healaon Pro, Neurive Inc., Gimhae, Republic of Korea) and ultrasound stimulation (ZenBud, NeurGear Inc., Rochester, NY, USA).

For E-taVNS, stimulation was applied bilaterally to the cymba conchae using conductive rubber electrodes. Parameters were set to 30 Hz at an intensity of level 5 (2.5 mA), and each stimulation block lasted 30 minutes. A continuous wind-like sound was played during both active and sham sessions to mask any device-related noise.

For U-taVNS, stimulation was applied to the right cymba conchae. The ultrasound protocol used a centre frequency of 5.3 MHz, a pulse repetition frequency of 41 Hz, a 50% duty cycle, a peak mechanical output of 0.081 W/cm^2^, and an average pressure amplitude of 1.03 MPa.

### Experimental design and procedures

Before the experiment, participants completed several questionnaires: the Beck Anxiety Inventory (BAI) (Beck et al. 1988), the Beck Depression Inventory (BDI) (Beck et al. 1996), and the Multidimensional Assessment of Interoceptive Awareness (MAIA-2) (Mehling et al. 2018).

Each participant completed two sessions (active and sham), scheduled at least five days apart, with stimulation order counterbalanced across participants. At the beginning of each session, participants completed an associative memory task consisting of an encoding phase, a distraction task, and a retrieval phase (Figure 1A). Following the first completion of the task, participants received 30 minutes of taVNS. Immediately after stimulation, they completed the associative memory task again in the post-stimulation phase. At the end of each session, participants completed a side-effects questionnaire assessing any mild adverse sensations (e.g., headache, neck tension, nausea, tingling, or muscle contractions).

**Figure 1.**
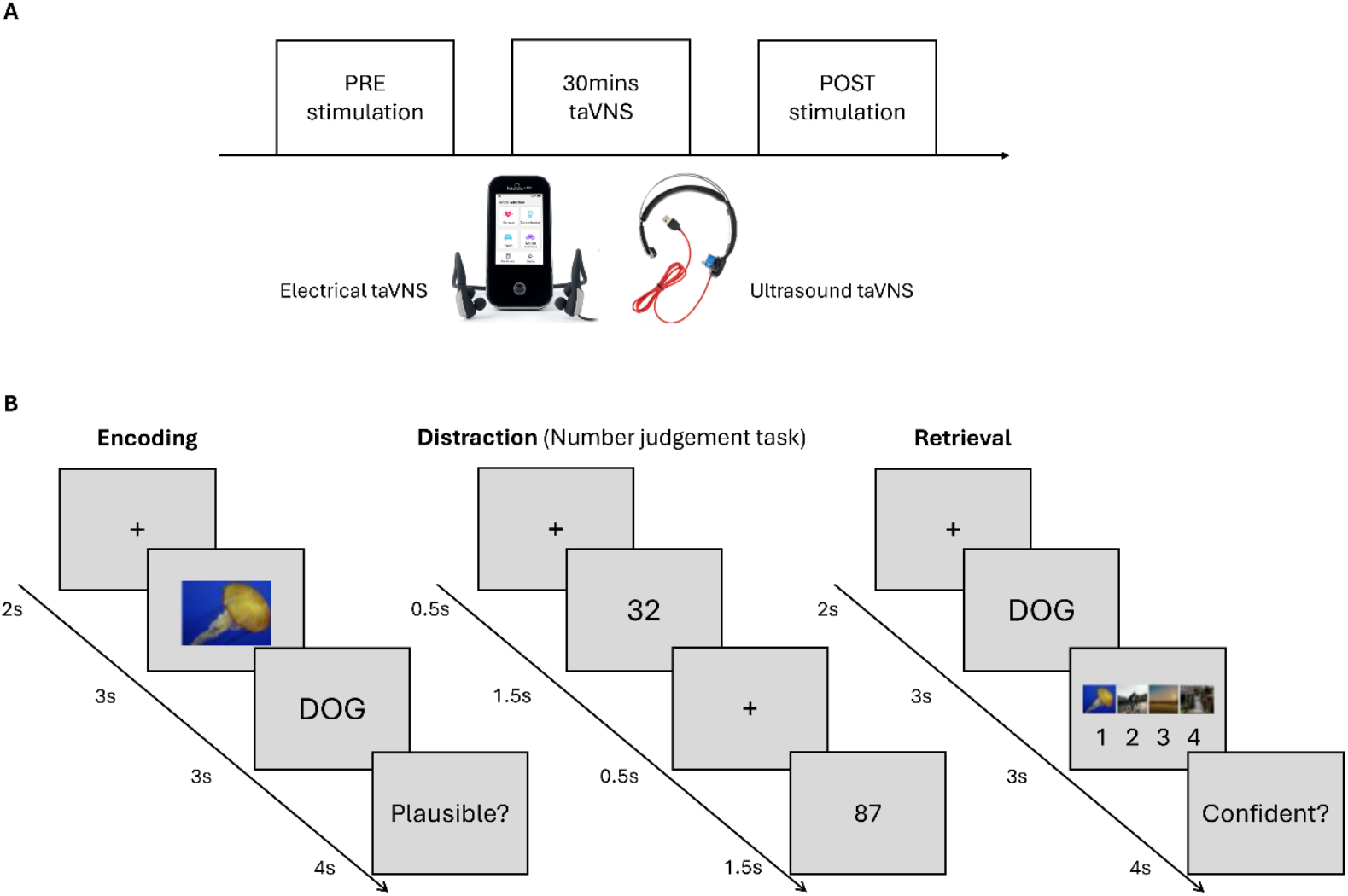
Experimental design and task procedure. (A) Overview of the stimulation protocol. Participants completed a pre-stimulation session, followed by 30 minutes of transcutaneous vagus nerve stimulation (taVNS), and a post-stimulation session. Two stimulation modalities were used: electrical taVNS and ultrasound taVNS. (B) Associative memory task. The task comprised encoding and retrieval phases separated by a distraction task. During encoding, a dynamic video was followed by a word, after which participants rated the plausibility of the video–word association on a 4-point scale. Each trial began with a 2s fixation. A number judgement distraction task (odd/even decision) was then administered. During retrieval, encoded words were presented and participants selected the associated video from four options, followed by a confidence rating (4-point scale).

### Associative memory task

We used the 4 dynamic video and 369 words from a prior study (Griffiths, Parish, et al. 2019). Figure 1B illustrates the task procedure. The associative memory task had two phases: encoding and retrieval. During the encoding phase, each trial presented a dynamic video for 3000ms in the middle of the screen, followed by a word that was shown on the screen for 3000ms. In this 6000ms, participants were instructed to vividly associate the video with the word to memory the video-word association.

Then, they were asked to rate plausibility of the video-word association on a 1-4 scale (1 = very implausible, 4 = very plausible) using Z X C V keys. Between the encoding and retrieval phases, the distraction task was administered. In number judgement task, a single number appeared at central fixation for 500ms, participants were required to judge whether the number was odd or even via designated buttons for only 1500ms. Stimuli were presented in randomized order. This task included 50 trials. During the retrieval phase, after each encoded word reappearing, participants had to select the associated video from the set of four videos stimuli, and then rated confidence (1 = not confident, 4 = very confident) using the Z X C V keys. There were strict time limits for retrieval phase, participants had 4000ms to select one of the four videos, followed by 4000ms to report their confidence. The associative memory task had 48 trials and each trial started with 2000ms fixation. Importantly, stimulus items differed across sessions to avoid repetition or carryover effects. The experiment was programmed and presented using PsychoPy (version 2024.2.4).

### Statistical analysis

Prior to conducting the central analyses, we cleaned the data using a series of exclusion criteria. First, trials where participants failed to choose a stimulus or provide a confidence rating were excluded (percentage of trials removed: 0.88%). Second, per-participant, reaction times for the memory responses which exceeded a ±2 SD threshold from the participant’s mean reaction time were excluded (percentage of trials removed: 4.87%). Third, we set out to remove participants with a memory performance less than 30%, reasoning that such performance is likely reflects chance performance (25%). Notably, however, no participant fell below this threshold, meaning all participants were included in the final analyses.

Our central analyses investigated the impact of tVNS on the accuracy and speed of episodic memory retrieval, focusing specifically on the responses made when selecting the target stimulus from the line-up of videos. We conducted analyses for electrical and ultrasound tVNS separately. We isolated the effects of active tVNS using difference scores, computed by subtracting performance following sham stimulation from performance under active conditions for each participant at each time point. We then conducted a series of one-tailed t-tests to evaluate our hypotheses that tVNS improves memory performance and hastens memory responses. First, we used one-sample t-tests to evaluate whether pre- and/or post-stimulation performance differed from zero (i.e., performance following sham stimulation). We then directly contrasted pre- and post-stimulation performance using paired-samples t-tests. For the reaction time analyses, we conducted a follow-up analysis to examine whether stimulation effects differed as a function of trial accuracy. We separated trials into correct and incorrect responses and repeated the planned contrasts described above for correct and incorrect responses separately.

To assess adverse effects, chi-square analyses were performed to compare the frequency of reported side effects between stimulation conditions (active vs. sham). Statistical significance was defined as *p* < 0.05.

## Results

### Demographic results

To confirm baseline comparability between groups, independent-samples t-tests and chi-square tests were performed on demographic and questionnaire scores (Table 1). No significant differences were observed between the e-taVNS and U-taVNS groups in sex distribution (χ^2^ = 0.00, p = 1.000) or age (t = −0.28, p = 0.781).

**Table 1.**
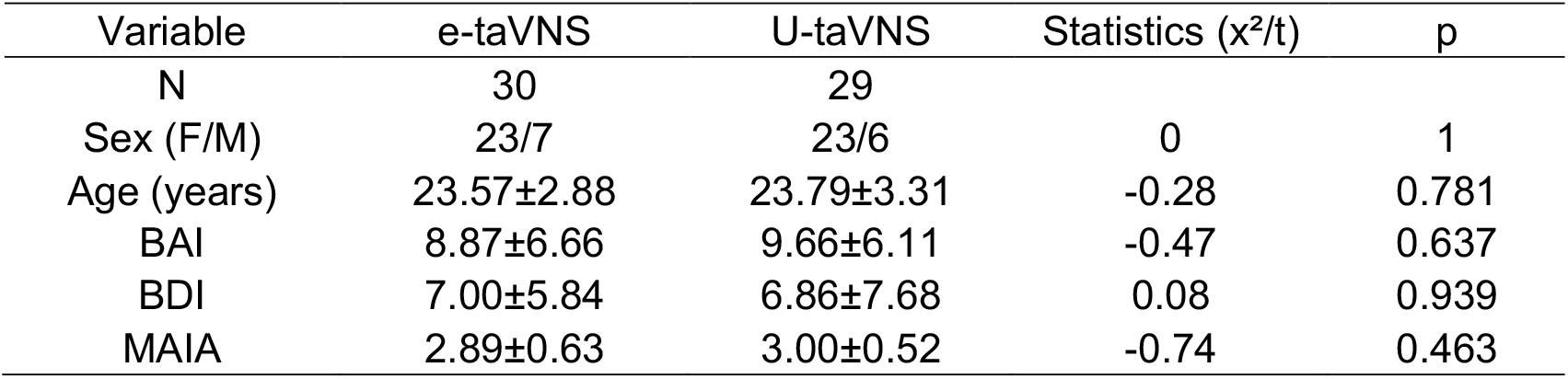
Participant demographic and characteristics.

| Variable | e-taVNS | U-taVNS | Statistics ($\chi^2/t$) | p |
| --- | --- | --- | --- | --- |
| N | 30 | 29 |  |  |
| Sex (F/M) | 23/7 | 23/6 | 0 | 1 |
| Age (years) | 23.57±2.88 | 23.79±3.31 | -0.28 | 0.781 |
| BAI | 8.87±6.66 | 9.66±6.11 | -0.47 | 0.637 |
| BDI | 7.00±5.84 | 6.86±7.68 | 0.08 | 0.939 |
| MAIA | 2.89±0.63 | 3.00±0.52 | -0.74 | 0.463 |

Likewise, the groups did not differ in anxiety (BAI: t = −0.47, p = 0.637), depression (BDI: t = 0.08, p = 0.939), or interoceptive awareness as assessed by the MAIA (t = −0.74, p = 0.463).

### Effects of taVNS on associative memory

Across conditions, participants on average were correct on 64.7% of trials (s.d.: 14.7%), with an average response time of 0.95 seconds (s.d.: 0.26s). We found no significant differences in memory performance or reaction times between the ultrasound and electrical tVNS groups [accuracy: t(57)=1.359, p=0.179; RT: t(57)=-1.698, p=0.095].

Neither electrical nor ultrasound tVNS had a significant impact on memory accuracy, with performance following active stimulation being no greater than sham stimulation [electrical: t(29) = 0.694, p = 0.247; ultrasound: t(27) = -0.747, p = 0.769; see Figure 2A] and no greater than performance prior to stimulation [electrical: t(29) = 0.004, p = 0.498; ultrasound: t(27) = -0.133, p = 0.552]. Bayesian analyses found anecdotal evidence for the null [electrical, active > sham: BF_01_ = 2.01; electrical, post > pre: BF_01_ = 2.56; ultrasound, active > sham: BF_01_ = 1.87; ultrasound, post > pre: BF_01_ = 2.42]. These results suggest that tVNS does not produce a significant enhancement in associative memory performance.

**Figure 2.**
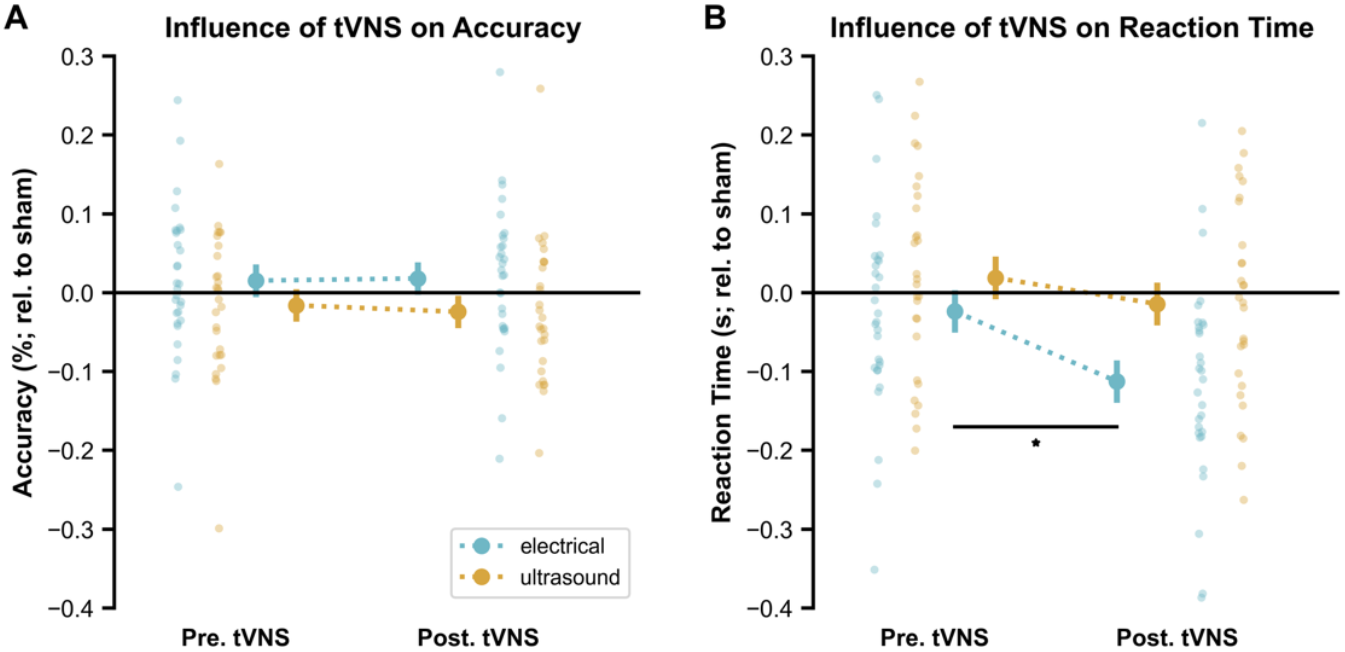
The influence of tVNS on behavioural performance during associative recall, relative to sham stimulation. **(A)** The impact of electrical tVNS (green) and ultrasound tVNS (yellow) on recall accuracy. Large dots in the centre of the plots represent the group, with accompanying errors describing standard deviation from the mean. Smaller dots on the sides of the plot depict individual participant effects. **(B)** The impact of electrical tVNS (green) and ultrasound tVNS (yellow), relative to sham stimulation, on response times. Plot details match Panel A. * p < 0.05.

In contrast, however, we did observe a substantial hastening of memory responses following electrical tVNS. Specifically, memory responses were faster following active electrical tVNS relative to sham stimulation [t(29) = -2.999, p = 0.003; see Figure 2B] and faster than responses prior to stimulation [t(29) = -1.887, p = 0.035]. This was a speeding of ∼100ms (or 8.5% change) relative to sham. While some may question whether the speeding of responses post-stimulation relative to pre-stimulation reflects a training effect, this idea is refuted by the similarly-hastened for active relative to sham stimulation as the “active vs. Sham” contrast compared conditions which incurred the same degree of training. The refutation of practice effects is reinforced by the absence of similar hastening effects following ultrasound tVNS [active < sham: t(27) = -0.391, p = 0.349, BF_01_ = 2.33; post < pre: t(27) = -0.694, p = 0.247, BF_01_ = 2.00]. Altogether, these results suggest that electrical tVNS hastens memory-related responses.

One critical open question remains: is the stimulation-induced hastening of responses something specific to successful associative memory retrieval, or is it a more general effect (e.g., hastening sensorimotor processing)? To answer this, we split trials based on retrieval success, reasoning that if tVNS aids memory retrieval, then these effects should only be observable for successfully retrieved items. In contrast, if tVNS exerts a more general sensorimotor effect, it should be observable regardless of memory retrieval. When analysing recalled items, we observed a significant hastening of reaction times for electrical tVNS relative to sham and pre-stimulation measurements [active vs. sham: t(28) = -2.816, p = 0.004; post < pre: t(28) = -1.907, p = 0.033; see Figure 3B]. For forgotten items, though there was a similar hastening for active relative to sham stimulation [t(27) = -1.766, p = 0.044], there was no hastening relative to pre-stimulation [t(27) = -0.357, p = 0.362]. Given that analysis of responses times between active and sham sessions, prior to stimulation, revealed a similar, trending hastening for forgotten items [t(27) = -1.486, p = 0.074], it would suggest that faster responses for forgotten items reflects a general hastening from sham to active sessions, rather than an effect that is specific to stimulation. Therefore, these results suggest that tVNS results in a hastening of responses times that is tied to successful memory retrieval and is not reflective of a more hastening of general cognitive/sensorimotor processes.

**Figure 3.**
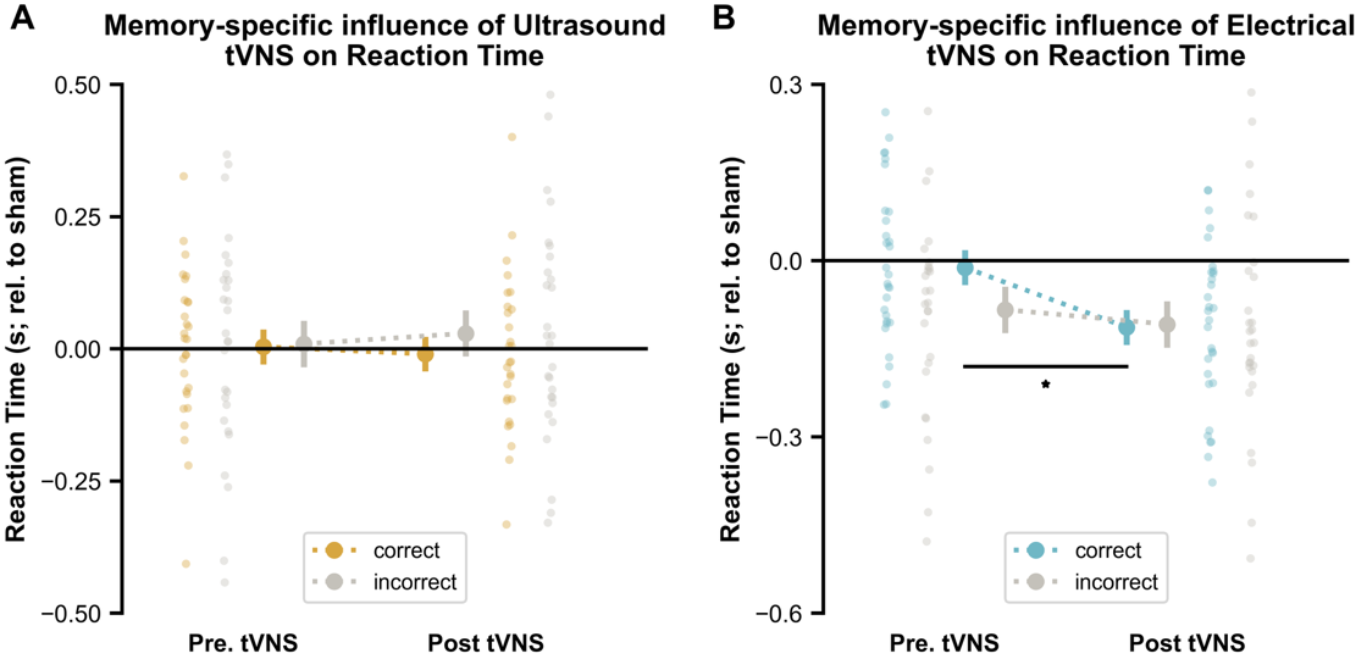
The influence of tVNS on reaction times as a function of recall success, relative to sham stimulation. **(A)** The impact of ultrasound tVNS on reaction time for correctly (coloured) and incorrect (gray) memory responses. Large dots in the centre of the plots represent the group, with accompanying errors describing standard deviation from the mean. Smaller dots on the sides of the plot depict individual participant effects. **(B)** The impact of electrical tVNS on response times as a function of recall success. Plot details match Panel A. * p < 0.05.

### Aversive effects results

A chi-square analysis was conducted to assess whether the incidence of aversive effects differed by stimulation modality (Table 2). Overall, no significant aversive effects were observed relative to sham in either modality. However, skin irritation was more frequently reported in the E-VNS group than in the U-VNS group, yielding a significant association between stimulation type and skin irritation (χ^2^ = 3.86, p = .049). This finding indicates that U-taVNS was associated with a lower incidence of skin irritation compared with E-taVNS.

**Table 2.** The summary of aversive effects. Values shown as n. * p < 0.05.

|  | Electrical taVNS |  |  |  | Ultrasound taVNS |  |  |  | E-taVNS vs U-taVNS |  |
| --- | --- | --- | --- | --- | --- | --- | --- | --- | --- | --- |
| | Active | Sham | $\chi^2$ | p | Active | Sham | $\chi^2$ | p | $\chi^2$ | p |
| Skin irritation | 6 | 3 | 1.20 | 0.273 | 1 | 0 | 1.02 | 0.313 | 3.86 | 0.049* |
| Headache | 1 | 2 | 0.35 | 0.552 | 0 | 0 | - | - | 0.98 | 0.321 |
| dizziness | 0 | 2 | 2.08 | 0.150 | 1 | 0 | 1.02 | 0.313 | 1.05 | 0.305 |
| nausea or vomiting | 0 | 1 | 1.02 | 0.313 | 0 | 0 | - | - | - | - |
| facial drooping | 0 | 0 | - | - | 0 | 0 | - | - | - | - |
| hoarse voice | 1 | 1 | 0.00 | 1.000 | 0 | 0 | - | - | 0.98 | 0.321 |
| pain or tingling in the throat or neck | 1 | 2 | 0.35 | 0.552 | 0 | 0 | - | - | 0.98 | 0.321 |
| cough | 1 | 0 | 1.02 | 0.313 | 0 | 0 | - | - | 0.98 | 0.321 |
| shortness of breath | 0 | 0 | - | - | 0 | 0 | - | - | 0.98 | 0.321 |
| Ear pain | 5 | 3 | 1.48 | 0.224 | 3 | 3 | 0.00 | 1.000 | 0.5 | 0.478 |
| others | 3 | 3 | 0.00 | 1.000 | 1 | 0 | 1.02 | 0.313 | 1 | 0.317 |

## Discussion

There is growing interest in the therapeutic potential of taVNS in tackling memory issues across society. Here, we investigated its ability to enhance the cued recall of crossmodal associative memories, diverging from established approaches using recognition tests and unimodal memories. Contrary to recent studies of recognition memory, we did not find that taVNS enhanced cued recall accuracy. Intriguingly, however, we did observe a hastening of response times following taVNS that was specific to memories that were correctly recalled. This also contrasts with studies of recognition memory which have often not observed taVNS-driven effects on reaction time. When combining our results with the existing literature, it suggests that taVNS exerts widespread and diverse effects over human memory.

Our results contribute to a growing, nuanced understanding of the impact of taVNS on the accuracy of longer-term (i.e., beyond the timespan of working) memory.

Several studies have demonstrated that taVNS can enhance recognition performance for a variety of stimuli, including images (Ventura-Bort et al. 2021; Ludwig et al. 2025; Ventura-Bort et al. 2025), faces (Jacobs *et al*. 2015), and Kanji word pairs (Annaka et al. 2025). However, these effects are often conditional. For example, taVNS preferentially boosts recognition for negative emotional stimuli, with smaller effects for neutral stimuli (Ventura-Bort *et al*. 2021; Ludwig *et al*. 2025). Furthermore, the recognition-related benefits of taVNS often for recollected stimuli, but not those which are recognised through familiarity-based processes (Giraudier et al. 2020). Our observation that taVNS does not enhance memory when participants must actively recall a paired associate suggests that the nature of the retrieval process also moderates the effectiveness of taVNS in modulating memory. Notably, our results align with the observation that taVNS fails to influence the free recall of verbal stimuli (Mertens *et al*. 2020), suggesting that the absence of memory effects is not attributable to particular stimuli or ways of administering a recall test, but is instead better attributed to taVNS not influencing the act of active recall itself.

Intriguingly, however, we did observe an effect of electrical taVNS on reaction times when successfully recalling a memory. Several studies in other cognitive domains have demonstrated that taVNS can hasten response times, including auditory oddball detection (Gurtubay et al. 2023) and conflict processing (Fischer et al. 2018). While these results suggest that taVNS has a more general cognitive effect on response times, such effects cannot explain why we observe hastening of response times specifically for correctly recalled items. Instead, the hastening of responses times must be driven by something specific to memory trace itself. As we applied taVNS offline, prior to both the encoding and the retrieval stages, this could be related to how the memory is stored during encoding, facilitating access during subsequent retrieval, or could be due to a hastening of the memory search process during active recall. Future research could disentangle these ideas by either exploring whether our reaction time effects can continue to be observed after the effects of taVNS have washed out, which would favour an encoding-based account, or whether the effects are enhanced when taVNS is delivered immediately before/during retrieval, favouring a memory search account.

The neural mechanisms that underpin this memory-specific hastening of reaction times remains to be seen. A viable candidate is the formation and/or retrieval of associative links by the hippocampus. Indeed, a past study using the same paradigm used here have demonstrated that successful recall is predicted by hippocampal engagement during the formation and retrieval of the associative links between stimuli (Griffiths, Parish, *et al*. 2019). Given the impact of taVNS on hippocampal function (Kraus *et al*. 2007; Murphy *et al*. 2023; Rajiah *et al*. 2024), it seems plausible to suggest that taVNS may be aiding hastening the recall process by augmenting hippocampal activity at critical junctures during either encoding or retrieval. An alternative candidate mechanism is alpha oscillations. taVNS has been shown to desynchronise alpha oscillations (Sharon *et al*. 2021), and such alpha desynchronisation is known to facilitate memory formation and retrieval in the task we used here (Michelmann et al. 2016; Griffiths, Mayhew, et al. 2019). Future research which pairs taVNS with electrophysiology or neuroimaging may be key for delineating these theories.

However, we did not observe any effects of ultrasound taVNS on associative memory performance. This null result should be interpreted with caution due to two key differences between our ultrasound and electrical protocols. First, whereas our electrical taVNS was applied bilaterally, the ultrasound stimulation was unilateral.

Given that most conventional protocols target the left ear to mitigate right-sided cardiac risks, it is possible that bilateral ultrasound application is necessary to reach the threshold for measurable cognitive effects. Second, the two modalities operate via distinct biophysical mechanisms. While electrical stimulation triggers direct, current-induced depolarization, ultrasound utilizes mechanical pressure waves to deform neuronal membranes and gate mechanosensitive ion channels (Blackmore et al. 2019). These fundamental differences suggest that ultrasound may require specific parameter optimization to effectively modulate the vagal afferent pathways involved in associative memory.

Several limitations warrant consideration when interpreting these results. First, the sample size was relatively small for both stimulation modalities, potentially limiting the statistical power to detect subtle effects. Second, this study focused exclusively on acute outcomes; consequently, it remains unclear whether these effects persist or accumulate through repeated sessions. Third, the lack of concurrent physiological monitoring such as heart rate variability or pupillometry biomarkers, precludes a direct validation of the underlying mechanistic pathways.

In sum, while taVNS did not improve the accuracy of active recall, we find that electrical taVNS significantly hastens response times for correctly recalled items, suggesting that it may be a valuable tool for facilitating of memory retrieval processes.

## Author contributions

Conceptualization: BJG, JJ; Methodology: BJG, JJ, MK, HC, JS; Investigation: ZH, IC, JJ; Writing— BJG, JJ; Writing—review & editing: BJG, JJ, MK, HC, JS

